# *Ripk1* Deficiency or K376R Mutation in Dendritic Cells Promotes Potent Antitumor Immunity

**DOI:** 10.64898/2026.02.04.703775

**Authors:** Yangjing Ou, Lingxia Wang, Xixi Zhang, Xiaoming Zhao, Lin Mao, Xiaoxia Wu, Landian Hu, Xiangyin Kong, Haibing Zhang

**Affiliations:** Key Laboratory of Nutrition and Metabolism, Shanghai Institutes of Nutrition and Health, Chinese Academy of Sciences, University of Chinese Academy of Sciences, Shanghai 200031, China; CAS Key Laboratory of Tissue Microenvironment and Tumor, Shanghai Institute of Nutrition and Health, Chinese Academy of Sciences, Shanghai 200031, China

## Abstract

Dendritic cells (DCs) are critical for initiating adaptive immunity through antigen presentation and T cell priming. While immunogenic cell death (ICD) enhances anti-tumor immunity, how programmed cell death in DCs shapes tumor immune responses remains unclear. Receptor-interacting protein kinase 1 (RIPK1) regulates apoptosis, necroptosis, and inflammation. Here, we show that mice with disrupted RIPK1 ubiquitination on K376 (*Ripk1^fl/K376R^ Cd11c-Cre*, *Ripk1^DC-KO/K376R^*) develop inflammation and autoimmunity, phenocopying DC-specific RIPK1-deficient(*Ripk1^fl/fl^ Cd11c-Cre*, *Ripk1^DC-KO^*) mice. These phenotypes were rescued by the genetic ablation of *Ripk3* or *Mlkl*, establishing that RIPK1 ubiquitination is essential for suppressing DC necroptosis to maintain immune homeostasis. Remarkably, both *Ripk1^DC-KO^* and *Ripk1^DC-KO/K376R^* mice exhibit robust resistance to tumor growth, characterized by significantly enhanced T cell infiltration and activation. Mechanistically, we show that *Ripk1* deficiency or the loss of its K376 ubiquitination augments antigen-presenting capacity of DCs through both necroptosis-dependent and -independent pathways. Furthermore, adoptive transfer of T cells from *Ripk1^DC-KO^* mice further amplifies anti-tumor immunity in recipient hosts. These results identify RIPK1 as a critical immunological checkpoint in DCs that constrains their immunogenic potential. Targeting this pathway represents a promising therapeutic strategy to potentiate T cell-mediated cancer immunotherapy.

## INTRODUCTION

The tumor microenvironment (TME), composed of stromal, vascular, and immune cells, plays a critical role in shaping cancer progression and immune evasion^1^. Beyond intrinsic genetic alterations in tumor cells, extrinsic factors within the TME-particularly immune regulation-determine whether tumors are recognized or eliminated by the host immune system^2–4^. While chronic inflammation is a known driver of tumorigenesis in organs such as the liver and stomach^5^, emerging evidence suggests that carefully orchestrated inflammatory signals can also enhance anti-tumor immunity when appropriately controlled. Therefore, it still remains an effective approach to inhibit specific tumor invasion by inducing sufficient inflammation and minimizing inflammation toxicities^6–10^.

Programmed cell death (PCD), including apoptosis, pyroptosis and necroptosis, have been recognized not only for maintaining tissue homeostasis but also for modulating immune responses^11^. Necroptosis, a regulated form of lytic cell death mediated by RIPK1, RIPK3 and MLKL, has been proposed to promote inflammation and anti-tumor immunity through the release of danger-associated molecular patterns (DAMPs)^12,13^. The immunogenic potential of basophil or tumor cell necroptosis has been demonstrated in several experimental studies and clinical trials^14–16^, which primarily originated from immune responses rather than antigen characteristics^17^. But the functional relevance of necroptosis in immune cells, especially in dendritic cells (DCs), remains poorly understood.

DCs are central to initiating adaptive immune responses through antigen uptake, processing, and T cell priming. However, whether and how necroptosis in DCs modulates their immunogenicity and shapes anti-tumor responses remain unclear. RIPK1 serves as a critical molecular hub that integrates pro-survival and pro-death signals through context-dependent post-translational modifications. Complete loss of RIPK1 leads to perinatal lethality^18–21^, but specific deletion in select immune compartments is tolerated and has revealed tissue-specific regulatory functions^22^. Specifically, DC-specific deletion of RIPK1 has been shown to induce spontaneous inflammation and autoimmunity, highlighting its role in maintaining immune tolerance^22^. In addition, we and others have shown that ubiquitination of RIPK1 at lysine 376 (K376) is essential for suppressing apoptosis and necroptosis during development^18,19^, although its role in DC-mediated immunity remains unexplored.

Here, we showed that DC-specific deletion of RIPK1 or disruption of its ubiquitination at lysine 376 leads to similar systemic inflammation and autoimmunity, largely dependent on RIPK3- and MLKL-mediated necroptosis. Strikingly, both genotypes exhibit broad resistance to tumor growth, accompanied by enhanced T cell activation and effector responses. Mechanistically, we show that *Ripk1* deficiency or loss of RIPK1 ubiquitination reprograms DCs toward a hyper-immunogenic state through both necroptosis-dependent and -independent mechanisms. Adoptive transfer of T cells from *Ripk1*-deficient mice confers tumor resistance to naïve hosts, underscoring the critical role of DC-intrinsic RIPK1 in shaping T cell–mediated anti-tumor immunity. Together, our findings uncover a previously unrecognized function of RIPK1 in restraining the immunostimulatory capacity of DCs and establish RIPK1 as a potential target for reprogramming DCs to enhance cancer immunotherapy.

## RESULTS

### Mice with the RIPK1 K376R mutation in DCs develop inflammation and autoimmunity similar to DC-specific RIPK1 knockout mice

RIPK1 controls pro-survival and pro-death signaling through context-dependent post-translational modifications, including ubiquitination. In particular, ubiquitination at lysine 376 (K376) is critical for restraining both apoptosis and necroptosis^18,19^. To investigate the role of K376 ubiquitination in dendritic cells (DCs), we generated mice expressing the RIPK1 K376R mutant allele specifically in CD11c⁺ cells by crossing *Ripk1^fl/K376R^* mice with *CD11c-Cre* mice (hereafter *Ripk1^DC-KO/K376R^*). These mice survived to adulthood, and RIPK1 mRNA and protein levels in DCs were comparable to controls, indicating that the K376R mutation does not impair RIPK1 expression in DCs (**Figure S1A-C**).

We observed significantly reduced frequencies of conventional DCs (cDCs) and plasmacytoid DCs (pDCs) in the spleens of *Ripk1^DC-KO/K376R^*mice, resembling the phenotype of *Ripk1^DC-KO^* mice (**Figure 1A, B**). In vitro, RIPK1^K376R^ BMDCs were highly susceptible to both spontaneous and stimulus-induced cell death under apoptotic and necroptotic conditions, similar to *Ripk1*-deficient DCs **(Figure 1C, D**), consistent with other studies^22,23^. These data suggest that disruption of RIPK1 ubiquitination sensitizes DCs to both apoptosis and necroptosis.

**Figure 1.**
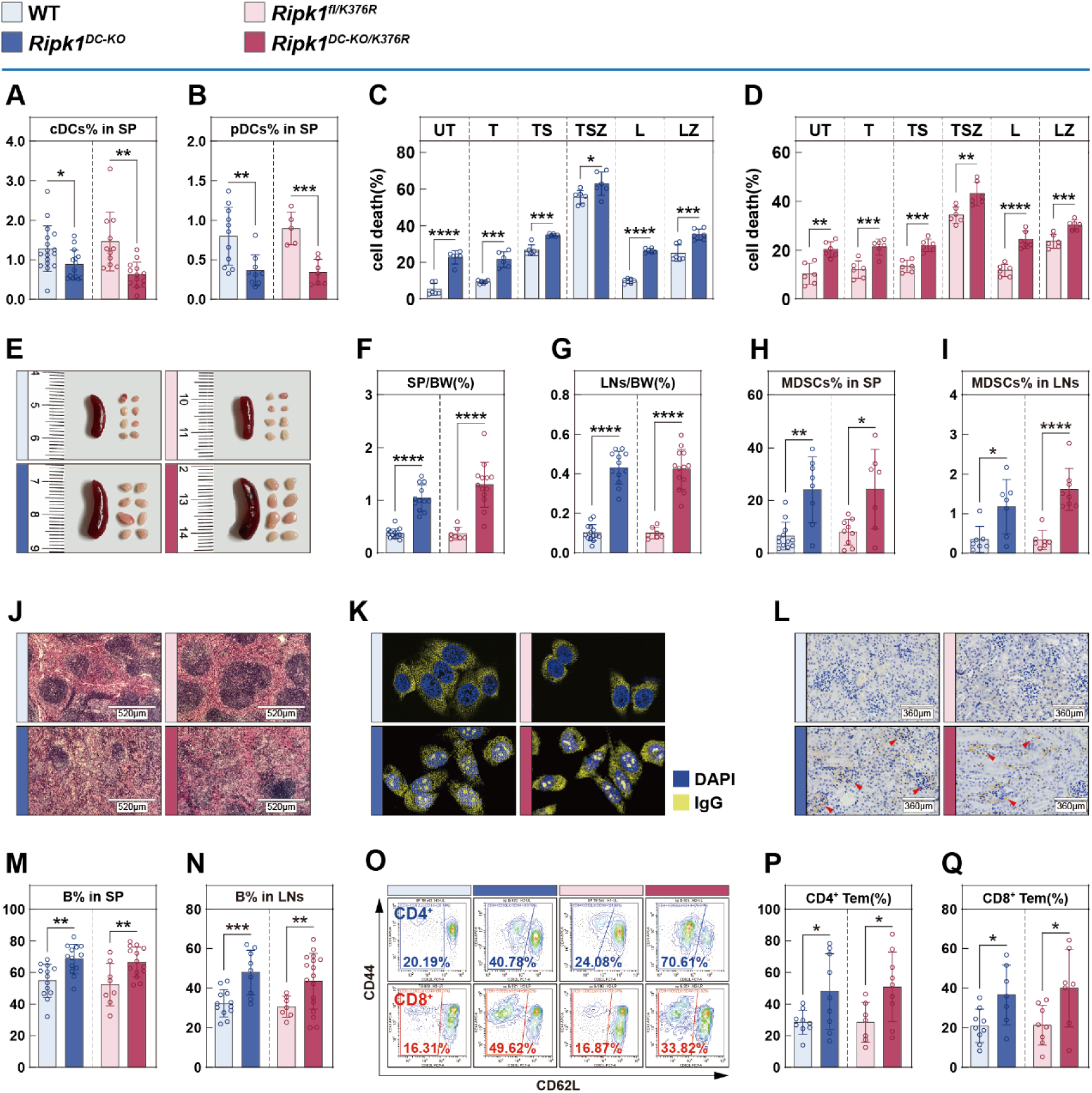
*Ripk1^DC-KO/K376R^* mice develop inflammation and autoimmunity. **(A, B)** Flow cytometric quantifications of splenic CD11c^+^ MHC-Ⅱ^+^ cDCs (A) and CD11c^lo^ CD317^+^ pDCs (B) in 6-week-old WT, *Ripk1^DC-KO^*, *Ripk1^fl/K376R^* and *Ripk1^DC-KO/K376R^* mice (n = 11-17). **(C, D)** CD11c^+^ BMDCs were generated from WT and *Ripk1^DC-KO^* (C), *Ripk1^fl/K376R^* and *Ripk1^DC-KO/K376R^* (D) mice (n = 6), and cell death assay was performed with following drugs for 10 h: T (TNF-α, 20 ng/μL), S (SMAC mimetics, 100 nM), Z (zVAD-fmk, 20 μM), L (LPS, 100 ng/mL). **(E-G)** Representative images (E) and weight index (organ weight/body weight, the same as below) of spleens (F) and lymph nodes (G) (from the cervical region, axillae of the forelimbs and inguinal regions from top to bottom, the same as below) in 6-week-old WT, *Ripk1^DC-KO^*, *Ripk1^fl/K376R^* and *Ripk1^DC-KO/K376R^*mice (n = 7-14). **(H-J)** Flow cytometry of CD11b^+^ LY6C/G^+^ MDSCs in spleens (H, n = 14-17) and lymph nodes (I, n = 7-12) in 6-week-old WT, *Ripk1^DC-KO^*, *Ripk1^fl/K376R^* and *Ripk1^DC-KO/K376R^* mice and representative images of HE staining of spleens (J). **(K, L)** Representative images of mouse serum IgG immunofluorescence (K) and renal IgG immunohistochemistry (L) of WT, *Ripk1^DC-KO^*, *Ripk1^fl/K376R^* and *Ripk1^DC-KO/K376R^*mice. **(M, N)** Flow cytometric quantifications of B cells in spleens (M) and lymph nodes (N) of 6-week-old WT, *Ripk1^DC-KO^*, *Ripk1^fl/K376R^* and *Ripk1^DC-KO/K376R^* mice (n = 7-12). **(O-Q)** Representative images (O) and quantifications (P, Q) of flow cytometry of splenic CD62L^lo^ CD44^hi^ T cells (n = 7-17, including CD4^+^/CD8^+^ subsets) in 6-week-old WT, *Ripk1^DC-KO^*, *Ripk1^fl/K376R^* and *Ripk1^DC-KO/K376R^* mice. Data are presented as the mean ± standard deviation (SD) from at least three independent experiments, with dots representing individual samples. Statistical significance (*P* values) was determined by a two-tailed unpaired Student’s t-test or Wilcoxon test (*p < 0.05, **p < 0.01, ***p < 0.001, ****p < 0.0001).

Consistent with impaired DC homeostasis, *Ripk1^DC-KO/K376R^* mice exhibited pronounced splenomegaly and lymphadenopathy (**Figure 1E-G**), along with expansion of myeloid-derived suppressor cells (MDSCs) (**Figure 1H, I**). Histological analyses revealed disrupted splenic architecture (**Figure 1J**), and total splenocyte and DC numbers were increased, likely due to compensatory lymphoid hyperplasia (**Figure S2A-E**).

Given the inflammatory phenotypes, we assessed autoimmune manifestations. Serum from *Ripk1^DC-KO/K376R^* and *Ripk1^DC-KO^*mice displayed nucleolar anti-nuclear antibody (ANA) staining patterns on HepG2 cells, suggestive of anti-ribosomal reactivity (**Figure 1K**). Immune complex deposition was also observed in glomeruli (**Figure 1L**), indicating the development of autoimmunity.

Furthermore, immunophenotyping revealed expansion of B cells in spleen and lymph nodes (**Figure 1M, N, Figure S2F, G**), while T cell subset distributions remained largely unchanged in both peripheral and thymic compartments (**Figure S2H-X**), indicating that the autoimmune features were not due to altered T cell development. However, both *Ripk1^DC-KO^* and *Ripk1^DC-KO/K376R^* mice exhibited elevated frequencies of central memory T cells (**Figure 1O-Q**), suggesting increased antigen experience or chronic stimulation. B cell activation markers were not significantly upregulated (**Figure S2Y**), indicating a selective immune dysregulation in both *Ripk1^DC-KO^* and *Ripk1^DC-KO/K376R^* mice.

Histopathological analysis of non-lymphoid tissues also revealed preserved architecture in liver, kidney, and lung, with only minor perturbations (e.g., reduced goblet cells in the duodenum) (**Figure S3A-C**), indicating that the inflammatory and autoimmune manifestations were largely confined to the immune system in both *Ripk1^DC-KO^*and Ripk1^DC-KO/K376R^ mice.

Together, these results demonstrate that DC-specific expression of RIPK1^K376R^ is sufficient to drive systemic inflammation and autoimmunity, closely mirroring the phenotype observed upon DC-specific deletion of RIPK1. These findings underscore the functional importance of RIPK1 ubiquitination in maintaining DC immune tolerance and tissue homeostasis.

### Dendritic Cell Necroptosis Drives Systemic Inflammation and Autoimmunity in *Ripk1^DC-KO/K376R^* Mice

Necroptosis is a proinflammatory form of programmed cell death that can contribute to immune activation and tissue pathology. Prior studies demonstrated that blocking necroptosis protects *Ripk1^DC-KO^* mice from inflammatory disease, suggesting a functional role for RIPK1 in preventing immunogenic DC death^22^. However, *in vivo* quantification of necroptosis remains technically challenging, and RIPK1 is not strictly required for all necroptotic pathways^24^. To directly assess the contribution of necroptosis to the inflammatory phenotypes driven by RIPK1 ubiquitination disruption, we genetically ablated *Ripk3* or *Mlkl* in *Ripk1^DC-KO/K376R^* mice.

Restoration of splenic DC subsets in these compound mutants to near wild-type (WT) levels indicated that necroptosis is the principal driver of DC depletion in *Ripk1^DC-KO/K376R^* mice (**Figure 2A, B**). Importantly, genetic blockade of necroptosis also alleviated hallmark signs of systemic inflammation, including splenomegaly and lymphadenopathy, as shown by reductions in organ-to-body weight ratios and total immune cell counts (**Figure 2C-G**). Histological analyses confirmed normalization of splenic architecture, and flow cytometry revealed partial correction of MDSC dysregulation, particularly in lymphoid and myeloid compartments (**Figure 2H-J**).

**Figure 2.**
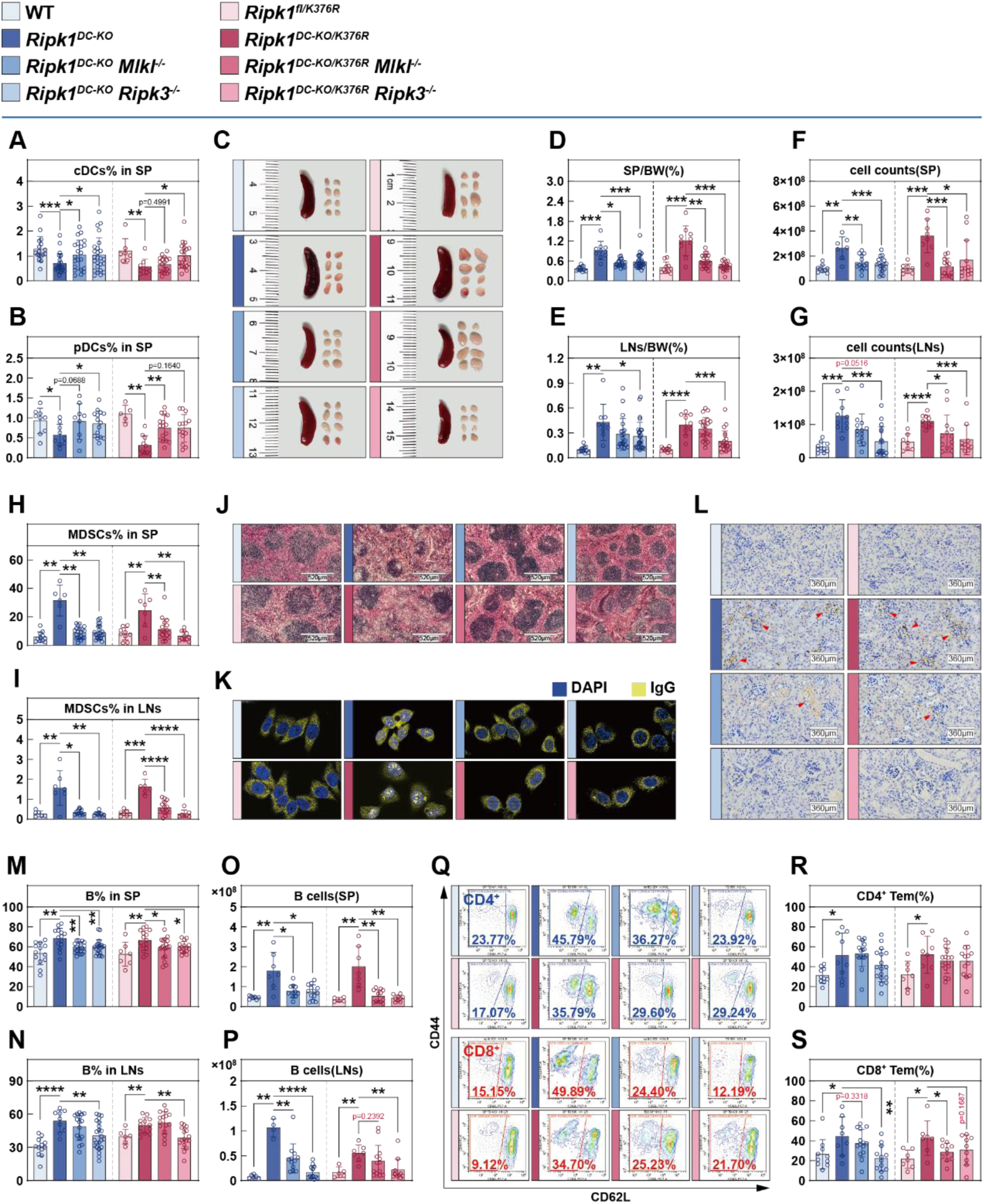
Dendritic cell necroptosis contributes to inflammation and autoimmunity in *Ripk1^DC-KO/K376R^* Mice. **(A, B)** Flow cytometric quantifications of splenic CD11c^+^ MHC-Ⅱ^+^ cDCs (A, n = 7-24) and CD11c^lo^ CD317^+^ pDCs (B, n = 5-16) in 6-week-old WT, *Ripk1^DC-KO^*, *Ripk1^DC-KO^ Mlkl^-/-^*, *Ripk1^DC-KO^ Ripk3^-/-^*, *Ripk1^fl/K376R^*, *Ripk1^DC-KO/K376R^*, *Ripk1^DC-KO/K376R^ Mlkl^-/-^* and *Ripk1^DC-KO/K376R^ Ripk3^-/-^* mice (n=11-17). **(E-G)** Representative images (C), weight index (D-E, n = 9-28) and cell counts (F-G, n = 7-17) of spleens and lymph nodes in 6-week-old WT, *Ripk1^DC-KO^*, *Ripk1^DC-KO^ Mlkl^-/-^*, *Ripk1^DC-KO^ Ripk3^-/-^*, *Ripk1^fl/K376R^*, *Ripk1^DC-KO/K376R^*, *Ripk1^DC-KO/K376R^ Mlkl^-/-^* and *Ripk1^DC-KO/K376R^ Ripk3^-/-^* mice. **(H-J)** Flow cytometry of CD11b^+^ LY6C/G^+^ MDSCs in spleens (H, n = 6-24) and lymph nodes (I, n = 6-14) in 6-week-old WT, *Ripk1^DC-KO^*, *Ripk1^DC-KO^ Mlkl^-/-^*, *Ripk1^DC-KO^ Ripk3^-/-^*, *Ripk1^fl/K376R^*, *Ripk1^DC-KO/K376R^*, *Ripk1^DC-KO/K376R^ Mlkl^-/-^* and *Ripk1^DC-KO/K376R^ Ripk3^-/-^* mice, and representative images of HE staining of spleens (J). **(K, L)** Representative images of mouse serum IgG immunofluorescence (K) and renal IgG immunohistochemistry (L) of WT, *Ripk1^DC-KO^*, *Ripk1^DC-KO^ Mlkl^-/-^*, *Ripk1^DC-KO^ Ripk3^-/-^*, *Ripk1^fl/K376R^*, *Ripk1^DC-KO/K376R^*, *Ripk1^DC-KO/K376R^ Mlkl^-/-^* and *Ripk1^DC-KO/K376R^ Ripk3^-/-^* mice. **(M-P)** Flow cytometric quantifications and cell count of B cells in spleens (M-O, n = 5-21) and lymph nodes (N-P, n =4-23) of 6-week-old WT, *Ripk1^DC-KO^*, *Ripk1^DC-KO^ Mlkl^-/-^*, *Ripk1^DC-KO^ Ripk3^-/-^*, *Ripk1^fl/K376R^*, *Ripk1^DC-KO/K376R^*, *Ripk1^DC-KO/K376R^ Mlkl^-/-^* and *Ripk1^DC-KO/K376R^ Ripk3^-/-^* mice. **(Q-S)** Representative images (Q) and quantifications (R, S) of flow cytometry of splenic CD62L^lo^ CD44^hi^ T cells (n = 6-17, including CD4^+^/CD8^+^ subsets) in 6-week-old WT, *Ripk1^DC-KO^*, *Ripk1^DC-KO^ Mlkl^-/-^*, *Ripk1^DC-KO^ Ripk3^-/-^*, *Ripk1^fl/K376R^*, *Ripk1^DC-KO/K376R^*, *Ripk1^DC-KO/K376R^ Mlkl^-/-^* and *Ripk1^DC-KO/K376R^ Ripk3^-/-^* mice. Data are presented as the mean ± SD from at least three independent experiments, with dots representing individual samples. Statistical significance (*P* values) was determined by a two-tailed unpaired Student’s t-test or Wilcoxon test (*p < 0.05, **p < 0.01, ***p < 0.001, ****p < 0.0001).

Necroptosis has been associated with autoimmune disorders in humans^25–27^. Notably, elevated MLKL expression is observed in peripheral blood mononuclear cells (PBMCs) from patients with systemic lupus erythematosus^28^. To determine whether DC necroptosis contributes to autoantibody production in our model, we measured circulating levels of anti-nuclear and anti-dsDNA antibodies. Remarkably, deletion of *Ripk3* or *Mlkl* significantly reduced autoantibody titers in *Ripk1^DC-KO/K376R^* mice (**Figure 2K**), indicating that DC necroptosis is a key driver of systemic autoimmunity in this context. Reduced immune complex deposition was also observed in glomeruli (**Figure 2L**), indicating the contribution of DC necroptosis in the development of systemic autoimmunity.

In addition, B cell expansion and other immune cell dysregulations recovered in spleen and lymph nodes (**Figure 2M-P, Figure S4A**-**N**), while the frequencies of CD4⁺ and CD8⁺ central memory T cells did not decline significantly, a consistent downward trend was observed in multiple individuals (**Figure 2Q-S**), suggesting that DC necroptosis contributes to, but does not fully account for, T cell activation. This implicates additional mechanisms or concurrent forms of immunogenic cell death in sustaining the autoimmune phenotype.

Together, these data establish that necroptotic death of DCs underlies the systemic inflammation and autoimmunity in *Ripk1^DC-KO/K376R^* mice, recapitulating the phenotypic features observed in *Ripk1^DC-KO^* mice.

### *Ripk1*-Deficient Dendritic Cells Promote Potent Antitumor Immunity

Inflammation and tumorigenesis are reciprocally linked with inflammation shaping the tumor microenvironment (TME) and influencing immune responses, stromal cell behavior, and tumor progression^29–31^. Given the systemic inflammation and immune activation observed in both *Ripk1^DC-KO^*and *Ripk1^DC-KO/K376R^* mice, we asked whether these DCs were sufficient to elicit antitumor immunity *in vivo*.

To test this, we intravenously injected *Ripk1^DC-KO^* and control mice with B16-F10 melanoma cells, a poorly immunogenic tumor model resistant to immune checkpoint blockade^32^. Two weeks after injection, *Ripk1^DC-KO^* mice showed markedly reduced lung metastases (by 72.4 ± 13.6%) and lower lung-to-body weight ratios (by 54.9 ± 19.4%) compared to controls (**Figure 3A-C**), indicating a robust antitumor response.

**Figure 3.**
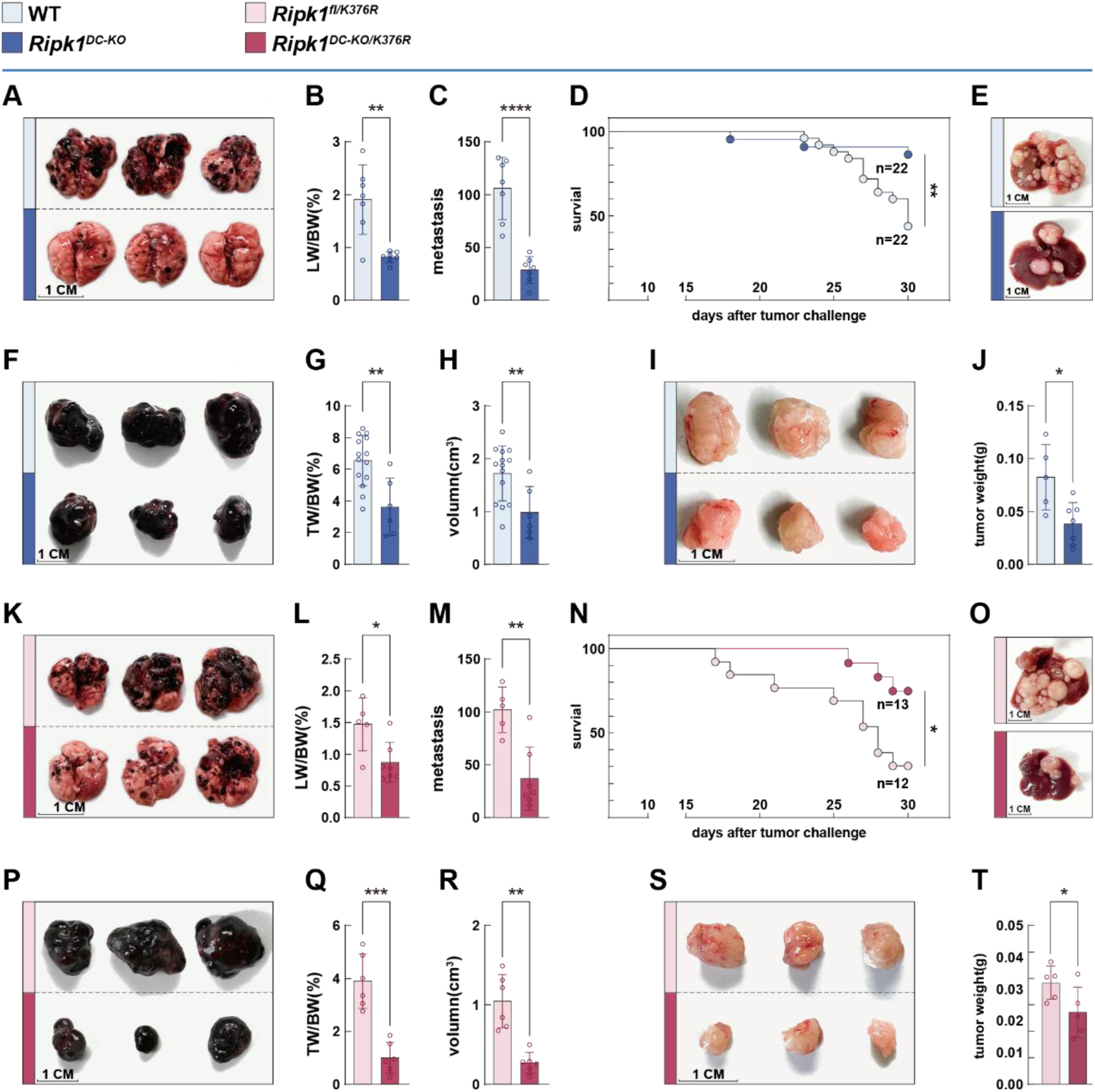
*Ripk1*-deficient and RIPK1^K376R^ DCs orchestrate pan-tumor resistance in mice. **(A–C)** Representative images of lung metastatic nodules (A), quantitative analysis of metastatic nodule counts per lung (B), and lung index (C) in WT and *Ripk1^DC-KO^* mice at 21 days post intravenous injection of B16-F10 melanoma cells (n = 7). **(D, E)** Kaplan-Meier survival curves (D) of WT and *Ripk1^DC-KO^*mice implanted with MC38 tumor cells (n = 22); representative images of liver metastatic lesions (E) collected from the above-mentioned mouse models at 30 days post tumor cell implantation. **(F–H)** Representative images of subcutaneous melanoma growth (F), quantitative analysis of tumor index (G), and tumor volumes (H) in WT and *Ripk1^DC-KO^*mice (n = 6–14). **(I, J)** Representative images of subcutaneous Pan02 pancreatic carcinoma growth (I) and quantitative statistics of tumor weight (J) in WT and *Ripk1^DC-KO^*mice (n = 5–9). **(K-M)** Representative images of lung metastatic nodules (K), quantitative analysis of metastatic nodule counts per lung (L), and lung index (M) in *Ripk1^fl/K376R^* and *Ripk1^DC-KO/K376R^* mice at 21 days post intravenous injection of B16-F10 melanoma cell (n = 7). **(N, O)** Kaplan-Meier survival curves (N) of *Ripk1^fl/K376R^* and *Ripk1^DC-KO/K376R^* mice implanted with MC38 tumor cells (n = 11-12); representative images of liver metastatic lesions (O) collected from the above-mentioned mouse models at 30 days post tumor cell implantation. **(P-R)** Representative images of subcutaneous melanoma growth (P), quantitative analysis of tumor index (Q), and tumor volumes (R) in *Ripk1^fl/K376R^* and *Ripk1^DC-KO/K376R^*mice (n = 6). **(S, T)** Representative images of subcutaneous Pan02 pancreatic carcinoma growth (S) and quantitative statistics of tumor weight (T) in *Ripk1^fl/K376R^* and *Ripk1^DC-KO/K376R^* mice (n = 5). Data are presented as the mean ± SD from at least three independent experiments, with dots representing individual samples. Statistical significance (*P* values) was determined by a two-tailed unpaired Student’s t-test or Wilcoxon test (*p < 0.05, **p < 0.01, ***p < 0.001, ****p < 0.0001).

To further assess whether *Ripk1*-deficient DCs could confer protection against other tumor types, we employed a transient intra-splenic inoculation model using MC38 colorectal carcinoma cells, followed by splenectomy. Notably, *Ripk1^DC-KO^* mice exhibited improved overall survival and reduced metastatic burden over 30 days of observation (**Figure 3D**, **E**), suggesting that DC-specific *Ripk1* deficiency primes systemic anti-tumor immunity independent of continuous tumor antigen exposure.

We next evaluated tumor growth in a subcutaneous implantation model. B16-F10 tumor growth was significantly delayed in *Ripk1^DC-KO^* mice, as indicated by reduced tumor weight (by 44.9 ± 12.4%) and volume (by 42.8 ± 14.6%) two weeks post-implantation (**Figure 3F-H**). Similarly, growth of a pancreatic tumor model (Pan02) was also suppressed in these mice 40 days after implantation (**Figure 3I, J**).

Given the immunosuppressive nature of the hepatic environment and its high susceptibility to metastatic colonization, we reasoned that spontaneous tumor control in the liver would indicate a potent, antigen-independent immune response. Indeed, *Ripk1*-deficient DCs mounted significant antitumor effects even in the liver, highlighting their capacity to break immune tolerance and initiate tumor rejection.

Together, these data demonstrate that DC-specific *Ripk1* deficiency confers a strong and broad-spectrum antitumor response across multiple tumor models and anatomical sites, likely through inflammation-driven immune reprogramming.

### RIPK1 K376R Mutation in Dendritic Cells Enhances Antitumor Immunity Similar to *Ripk1* Deficiency

Although the immune activation elicited by RIPK1 K376R mutation in dendritic cells (DCs) phenocopies that of *Ripk1*-deficient DCs, it remains unclear whether this mutation is sufficient to recapitulate the robust antitumor immunity observed in *Ripk1^DC-KO/K376R^* mice.

To address this, we first evaluated lung metastasis following intravenous injection of B16-F10 melanoma cells, a poorly immunogenic and checkpoint-resistant tumor model^32^. By day 21 post-injection, *Ripk1^DC-KO/K376R^* mice exhibited a marked reduction in metastatic nodules in the lungs, as well as significantly lower lung-to-body weight ratios compared to WT controls (**Figure 3K-M**). These findings suggest that the RIPK1 K376R mutation in DCs effectively impairs melanoma colonization and outgrowth in the lung.

We next employed the MC38 colorectal cancer liver metastasis model to assess broader antitumor efficacy. In this setting, *Ripk1^DC-KO/K376R^* mice demonstrated enhanced overall survival and a substantial decrease in hepatic metastatic lesions at 30 days post-inoculation ((**Figure 3N, O**).

To further assess the impact on tumor progression in primary tissues, we conducted subcutaneous implantation of both B16-F10 melanoma and pancreatic tumor cells. In both models, *Ripk1^DC-KO/K376R^* mice showed significantly delayed tumor growth and reduced tumor size and weight relative to controls ((**Figure 3P-T**). These effects were observed in the absence of exogenous adjuvants or engineered tumor antigens.

Taken together, these data demonstrate that disruption of RIPK1 K376 ubiquitination in DCs is sufficient to induce potent antitumor immunity across diverse tumor types and anatomical sites. The antitumor effect appears to be mediated through immune system reprogramming rather than antigen-specific responses, consistent with the immunostimulatory consequences of dysregulated cell death and inflammation in *Ripk1DC-KO/K376R* mice.

### RIPK1 Deletion or K376R Mutation in Dendritic Cells Shapes T Cell Responses to Promote Antitumor Immunity

The induction of robust and tumor-specific T cell responses is essential for effective antitumor immunity. Given the enhanced tumor resistance and increased central memory T cell populations observed in *Ripk1^DC-KO^*and *Ripk1^DC-KO/K376R^* mice, we hypothesized that these mice may harbor functionally superior cytotoxic T lymphocytes (CTLs). Flow cytometric analyses revealed a significant increase in the frequency of CD8⁺ T cells co-expressing granzyme B and TNF in both spleens and tumor-draining lymph nodes (tdLNs) of mutant mice compared to WT controls (**Figure 4A-D**). This enrichment of cytotoxic effectors was accompanied by a slight increase in overall T cell abundance and a concomitant decrease in B cell frequency in tdLNs (**Figure 4E-H**), but the subsets of T cells in WT, *Ripk1^DC-KO^* and *Ripk1^DC-KO/K376R^* mice fellow the same general trend (**Figure S5A-D**). These findings indicate the characteristic functions of T cells in *Ripk1^DC-KO^*and *Ripk1^DC-KO/K376R^* mice, different from the pattern that mirrors previously reported observations where B cell-targeted immune checkpoint blockade (e.g., TIM-1) suppressed tumor growth by reprogramming the lymphoid environment in WT mice^33^.

**Figure 4.**
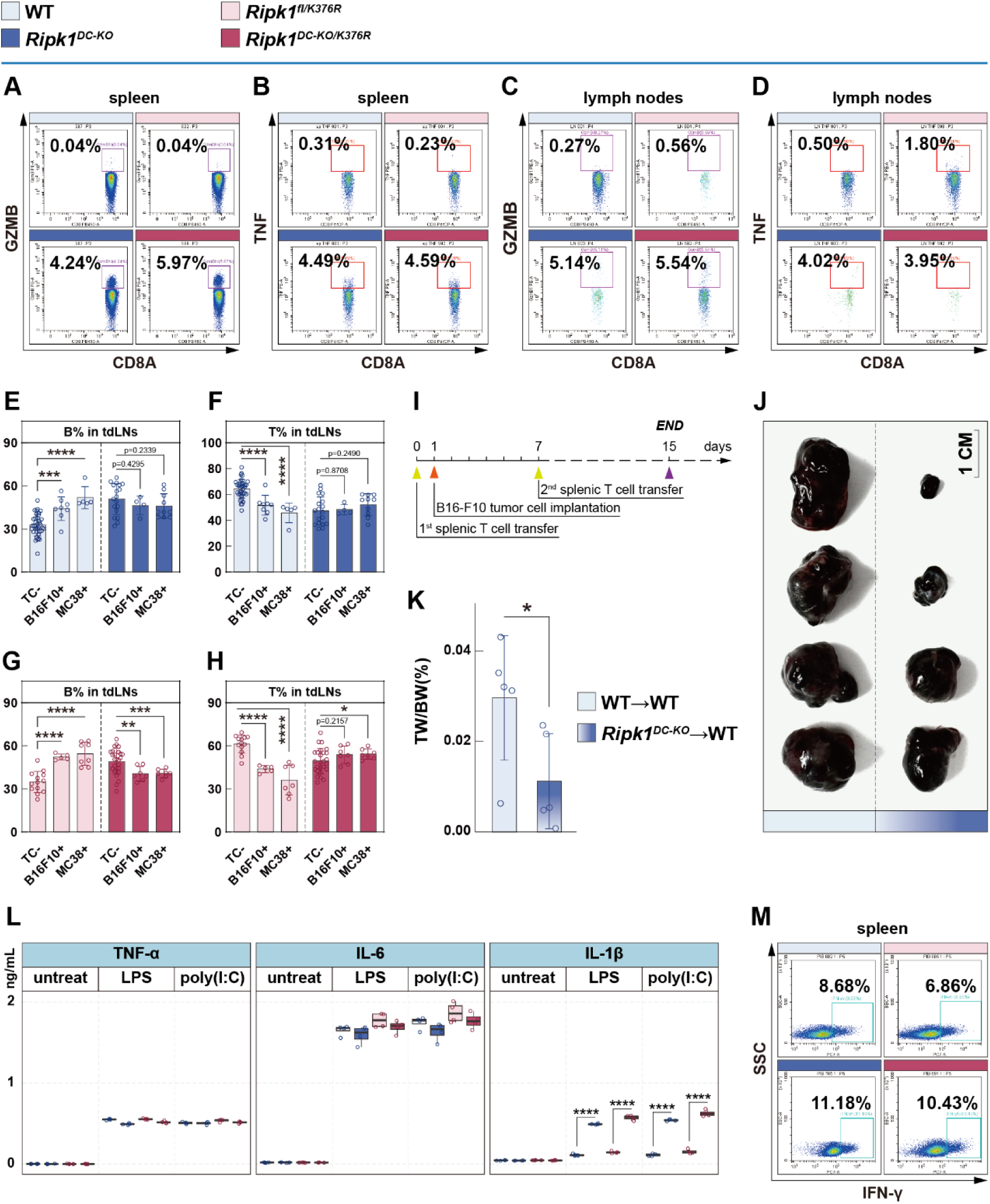
RIPK1 deletion or K376R mutation in dendritic cells promotes T cell activation and enhances antitumor immunity. **(A-D)** Representative images of flow cytometry of CD8^+^ GZMB^hi^ and CD8^+^ TNF^hi^ T cells in spleens (A, B) and lymph nodes (C, D) of 6-week-old mice of WT, *Ripk1^DC-KO^*, *Ripk1^fl/K376R^* and *Ripk1^DC-KO/K376R^* mice (tumor-challenged). **(E-H)** Flow cytometric quantifications of B cells and T cells in tumor-draining lymph nodes (n = 5-38). TC-: non-tumor-challenged. For MC38 tumor model, draining lymph nodes refer to hilar/periportal lymph nodes; for B16F10 subcutaneous tumor model, draining lymph nodes refer to right axillary lymph nodes (the same as below). **(I-K)** T cell transfer strategy (I) and representative images (J) and weight index (K) of B16-F10 subcutaneous tumors (n = 5). The transferred T cells were isolated from non-tumor-challenged WT and *Ripk1^DC-KO^* mice, enriched using EasySep™ Mouse T Cell Isolation Kit (STEMCELL, 19851) under sterile conditions, with 8×10⁴ cells injected each mouse each time. **(L)** ELISA detected cytokines of cultured medium of BMDCs treated with LPS (100 ng/mL) and Poly(I:C) (10 μg/mL) for 10 hours (n = 4). **(M)** Representative images of flow cytometry analysis of CD8^+^ IFN-γ^hi^ T cells in spleens of 6-week-old mice of WT, *Ripk1^DC-KO^*, *Ripk1^fl/K376R^* and *Ripk1^DC-KO/K376R^* mice. Splenic cells were isolated from non-tumor-bearing mice and stimulated with 10 ng/mL PMA (Sigma-Aldrich, P8139) and 1 μg/mL ionomycin (Sigma-Aldrich, N6386) for 4 h; 10 μg/mL brefeldin A (Sigma-Aldrich, B5936) was added after 2 h to inhibit cytokine secretion. Data are presented as the mean ± SD from at least three independent experiments, with dots representing individual samples. Statistical significance (*P* values) was determined by a two-tailed unpaired Student’s t-test or Wilcoxon test (*p < 0.05, **p < 0.01, ***p < 0.001, ****p < 0.0001).

To functionally test whether T cells from *Ripk1^DC-KO^* mice could confer tumor resistance, we adoptively transferred splenic T cells into WT recipients. These transferred T cells significantly delayed subcutaneous B16-F10 tumor growth (**Figure 4I-K**), though they failed to impact B16-F10 lung metastases (**Figure S5E**). Interestingly, in the MC38 liver metastasis model, immunofluorescence revealed increased T cell infiltration within tumor foci in recipient mice (**Figure S5F**), suggesting context-dependent T cell activity and tumor-site specificity.

Given the central role of dendritic cells (DCs) in orchestrating T cell responses, we next assessed the activation status of *Ripk1*-deficient and RIPK1^K376R^ DCs. Flow cytometry revealed comparable surface expression of canonical co-stimulatory molecules between mutant and WT DCs (**Figure S5G**), suggesting that differences in T cell priming were not driven by constitutive DC activation. To test their functional responsiveness, bone marrow-derived DCs (BMDCs) were stimulated with LPS or poly(I:C), followed by cytokine measurement. In the absence of stimulation, mutant DCs did not secrete elevated levels of inflammatory cytokines, including TNF-α (**Figure 4L**). Upon stimulation, TNF-α and IL-6 secretion remained comparable to WT levels, but IL-1β production was significantly enhanced in *Ripk1^DC-KO^* and *Ripk1^DC-KO/K376R^* DCs (**Figure 4L**). Furthermore, stimulation of splenic T cells from mutant mice led to greater IFN-γ production compared to controls (**Figure 4M**, **Figure S5H**), while T cells from lymph nodes showed no such difference (**Figure S5I**), possibly due to higher DC density in spleens.

Consistent with their proinflammatory profile, peripheral serum from mutant mice showed elevated levels of inflammatory cytokines (**Figure S5J**). Notably, deletion of *Mlkl* or *Ripk3* in these mice normalized serum cytokine levels (**Figure S5J**), implicating DC necroptosis as a major driver of systemic cytokine accumulation. Notably, IFN-γ levels remained persistently elevated in tumor-bearing *Ripk1^DC-KO^*mice compared to WT, even after tumor establishment, whereas other cytokines gradually returned to baseline (**Figure S5K, L**). This sustained IFN-γ signature may reflect a key mechanistic axis of antitumor immunity in this setting and warrants further investigation.

### Dendritic Cell Necroptosis Partially Mediates Antitumor Immunity Driven by *Ripk1* Deficiency or K376R Mutation

RIPK3-dependent necroptosis has been shown to contribute to antitumor effects in non-tumor cells, including myeloid populations, by promoting inflammatory cell death and enhancing immune activation^17^. Given our earlier findings that *Ripk1*-deficient and K376R mutant DCs confer broad antitumor resistance, we asked whether necroptosis is necessary for this phenotype.

To address this, we evaluated tumor responses in *Ripk1^DC-KO^* and *Ripk1^DC-KO/K376R^* with *Ripk3^−/−^*or *Mlkl^−/−^* mice using the same tumor models as previously described. In subcutaneous B16-F10 tumor implantation models, we observed that loss of Ripk3 or Mlkl significantly impaired the delayed tumor growth observed in both *Ripk1*-deficient or K376R mutant DCs mice (**Figure 5A, B**), suggesting a critical role for dendritic cell necroptosis in suppressing local tumor development. However, in contrast, the anti-metastatic capacity in the MC38 liver metastasis model remained largely intact (**Figure 5C, D**), indicating that DC necroptosis is dispensable for controlling metastatic tumor spread.

**Figure 5.**
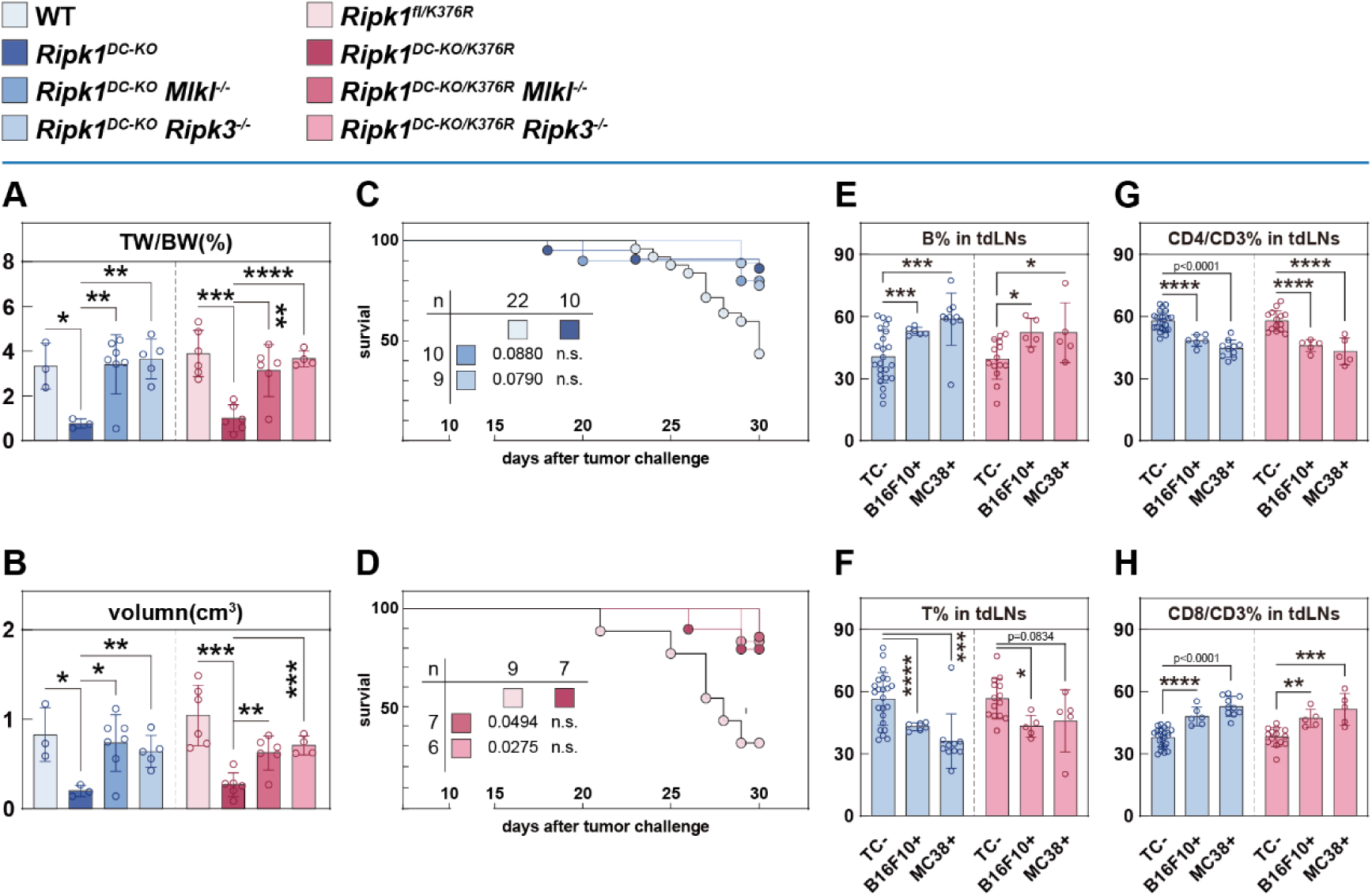
Dendritic cell necroptosis partially mediates the antitumor immunity induced by *Ripk1* deficiency or K376R mutation. **(A, B)** Weight index (A) and volume (B) of B16-F10 subcutaneous tumors in WT, *Ripk1^DC-KO^*, *Ripk1^DC-KO^ Mlkl^-/-^*, *Ripk1^DC-KO^ Ripk3^-/-^*, *Ripk1^fl/K376R^*, *Ripk1^DC-KO/K376R^*, *Ripk1^DC-KO/K376R^ Mlkl^-/-^* and *Ripk1^DC-KO/K376R^ Ripk3^-/-^* mice 14 days after implantation (n = 3-7). **(C, D)** Kaplan-Meier survival curves of WT, *Ripk1^DC-KO^*, *Ripk1^DC-KO^ Mlkl^-/-^*, *Ripk1^DC-KO^ Ripk3^-/-^*, *Ripk1^fl/K376R^*, *Ripk1^DC-KO/K376R^*, *Ripk1^DC-KO/K376R^ Mlkl^-/-^* and *Ripk1^DC-KO/K376R^ Ripk3^-/-^* mice implanted with MC38 tumor cells (n = 7-23). **(E-H)** Flow cytometric quantifications of B cell and T cells in tumor-draining lymph nodes (n = 6-24). Data are presented as the mean ± SD from at least three independent experiments, with dots representing individual samples. Statistical significance (*P* values) was determined by a two-tailed unpaired Student’s t-test or Wilcoxon test (*p < 0.05, **p < 0.01, ***p < 0.001, ****p < 0.0001).

In support of necroptosis-independent mechanisms, we observed that deletion of *Ripk3* or *Mlkl* did not significantly alter T cell subset distribution in the tumor-draining lymph nodes compared to WT controls (**Figure 5E-H**). This suggests that, while necroptosis contributes to local tumor control, sustained T cell activation and antitumor immunity in the metastatic context may be maintained through necroptosis-independent DC functions.

Together, these results demonstrate that dendritic cell necroptosis is a significant but non-exclusive driver of pan-tumor immune resistance conferred by RIPK1 deletion or K376R mutation, with additional downstream inflammatory and immunostimulatory pathways likely cooperating to shape antitumor immunity. A recent study indicated that distinct tumor cell death modes affected DCs differently, leading to varied tumor resistance^34^. Thus, we propose the tumor resistance from RIPK1^-/-^ or RIPK1^K376R^ is a product of combined regulation by multiple cell death modes—despite differential effects of DC death modes on tumor resistance—which still be beneficial.

## DISCUSSION

Our study uncovers a previously underappreciated role for RIPK1 ubiquitination in regulating dendritic cell (DC) death and antitumor immunity. Specifically, we demonstrate that DC-specific RIPK1 deletion or mutation at lysine 376 (K376R) induces robust resistance to both primary tumor growth and metastasis. While the suppression of necroptosis through *Ripk3* or *Mlkl* deletion significantly reversed the resistance to subcutaneous tumor growth, the antimetastatic immunity was surprisingly retained in both *Ripk1^DC-KO^* and *Ripk1^DC-KO/K376R^* mice. These findings genetically uncouple necroptosis execution from certain arms of antitumor defense and point toward alternative, necroptosis-independent mechanisms of immune activation.

Current research indicates that CD11c is not a reliable marker for the DC lineage, as transcription of the *Itgax* occurs in other myeloid-derived cells like macrophages. While this may not significantly conflict when identifying the specific defective sites causing systemic immune inflammation upon RIPK1 deletion in CD11c⁺ cells (which may include other cell types), it raises concerns regarding rigor in the context of tumor immunity. Therefore, however likely DCs are to be the direct cause of antitumor immunity, we cannot draw a definitive conclusion. It should be noted that no strictly conserved gene specific to DCs has been identified to date, and *Zbtb46*, once considered a candidate, was reported to be transcribed in splenic endothelial cells^35^. Although *Zbtb46* shows better conservation than *Itgax* among immune cells, the expression is restricted to conventional DCs and Langerhans cells, meaning it still cannot serve as a definitive pan-DC marker. Therefore, even though we acknowledge the limitations of the *CD11c-Cre* model, it still remains a common tool for studying DC characteristics and functions. Moreover, identifying the true source of tumor resistance in *CD11c-Cre* mice could help exclude irrelevant cell types, we consider it plausible that the broad population of CD11c⁺ cells, rather than DCs alone, may truly contribute to the strong antitumor phenotype observed in these mice.

Although DC necroptosis partially contributed to tumor control in localized tumor models, our data suggest that *Ripk1*-deficient and K376R mutant DCs can initiate antitumor immunity through mechanisms beyond necroptosis. Given the known plasticity of DC responses to environmental stimuli, it is plausible that alternative cell death pathways, such as apoptosis or pyroptosis, or basal DC activation, may also engage immunogenic signaling cascades in these settings^11^. Supporting this notion, we observed cleaved caspase activation and potential GSDMD processing in tumor-infiltrating DCs lacking RIPK1 or expressing the K376R mutant. Prior work has shown that sublethal pyroptotic signaling downstream of apoptotic caspases can contribute to antitumor immunity through the release of inflammatory mediators^11^. Additionally, recent evidence indicates that necrosome activation, even in the absence of full necroptotic execution, is sufficient to induce chemokine production from NF-κB primed hepatocytes^36^. We speculate that in the absence of MLKL or RIPK3, upstream necrosome signaling may persist and contribute to proinflammatory outputs without triggering full-blown lytic death, suggesting a mechanistic divergence between local and systemic tumor control in these models.

Besides, RIPK3-deficient macrophages exhibit altered lipid metabolism and increased PPAR activity, leading to M2 polarization and impaired tumor control in hepatocellular carcinoma models^37^, indicating the pleiotropic roles of RIPK3 complicate the interpretation of tumor outcomes following its deletion and may result in context-dependent immune modulation.

To further delineate how distinct death programs in DCs shape immunogenicity, we analyzed publicly available single-cell RNA-seq datasets of DCs under apoptotic or necroptotic stress (CD11c^hi^MHC-Ⅱ^hi^SIRP^+^ cells). Gene set variation analysis (GSVA) (**Figure S6A**) and PCA (**Figure S6B, C**) revealed that apoptotic signatures were more prevalent, but necroptosis-associated profiles correlated more strongly with immune-stimulatory gene expression (**Figure S6D, E**), including co-stimulatory molecules such as CD80, CD86, CD40, and MHC-II (**Figure S6F, G**). Importantly, these signatures originated from live cells, suggesting that sensitivity or engagement of death signaling pathways-not cell lysis itself-modulates DC activation state. Further enrichment analysis demonstrated that cells with high necroptosis scores exhibited enhanced expression of inflammatory and immune-related pathways compared to apoptosis-biased subsets (**Figure S6H**). These findings imply that the immunological impact of programmed cell death may be driven as much by sublethal signaling as by terminal cell demise, highlighting the immunomodulatory potential of death pathway engagement in DCs.

Recent studies have shown that nuclear RIPK1 can phosphorylate SMARCC2, a core component of the BAF (SWI/SNF) chromatin remodeling complex, thereby promoting nucleosome repositioning and inflammatory gene expression^40,41^. However, in RIPK1^K376R^ cells, enhanced RIPK1 phosphorylation paradoxically correlates with exacerbated inflammation, suggesting a disconnect between cytoplasmic activation and nuclear function^18,19^. We hypothesize that the absence of K376-linked ubiquitination impairs the interaction between RIPK1 and SMARCC2, as well as other nuclear partners, thereby abrogating nuclear RIPK1-mediated chromatin remodeling^41^. Moreover, BRG1 (SMARCA4), the ATPase subunit of the BAF complex, undergoes ubiquitin-mediated degradation upon CK1δ phosphorylation^42^. These observations raise the possibility that K376 ubiquitination facilitates chromatin association or complex stability, and its disruption compromises chromatin integrity, triggering an inflammatory form of cell death. Such a “chromatin-linked cell death” model might explain the heightened immunogenicity of K376R-mutant DCs^43^. However, we currently have limited understanding of whether these molecular events (aberrant chromatin compaction or decondensation, etc.) occur in the context of complete RIPK1 deletion.

In conclusion, our study delineates a scaffold-dependent, ubiquitination-regulated mechanism by which RIPK1 restrains DC cell death and inflammatory reprogramming. We show that necroptosis contributes partially to localized antitumor responses, but that resistance to metastasis persists even in the absence of this pathway, implicating alternative immune-stimulatory mechanisms. Furthermore, we propose that dysregulated nuclear functions of RIPK1 may underlie the intrinsic death sensitivity and immunogenicity of DCs in *Ripk1*-deficient settings. These findings advance our understanding of cell death signaling in innate immune cells and suggest that targeting scaffold functions of RIPK1 may enhance antitumor immunity without inducing overt lytic death.

## Materials and Methods

### Mice

All experimental mice were strictly maintained under specific pathogen–free (SPF) conditions (22–26 °C, 40–60% humidity, 12-h light/dark cycle) in the animal facility of Shanghai Institutes of Nutrition and Health, Chinese Academy of Sciences, University of Chinese Academy of Sciences. *Ripk1^K376R/+^*, *Ripk3^-/-^* and *Mlkl^-/-^* mice have been described in previous articles^19,44^. *CD11c-Cre* transgenic mice (Strain #: 008068) were purchased from The Jackson Laboratory (Bar Harbor, ME, USA). All strains were backcrossed to a pure C57BL/6 background for over 6 generations to ensure genetic homogeneity. All animal experimental protocols were reviewed and approved by the Institutional Animal Care and Use Committee (IACUC), and were strictly conducted in accordance with the guidelines for the Care and Use of Laboratory Animals issued by local regulatory authorities.

### Isolation and Culture of Murine BMDCs

Murine BMDCs were generated from 6–8-week-old mice. Mice were euthanized by CO_2_ inhalation; femurs and tibias were dissected, and surrounding skin/muscle tissues were removed. Epiphyses were cut off, and bone marrow cells were flushed out with RPMI 1640 medium (Gibco). The cell suspension was pipetted to disperse clumps. The pellet was resuspended in complete RPMI 1640 medium supplemented with 10% FBS (Bioind, 04-001-1A), 1% penicillin/streptomycin (Gibco, 15140122), 50 ng/mL IL-4 (PeproTech, 214-14-20), and 50 ng/mL GM-CSF (PeproTech, 315-03-50UG). Cells were seeded into 10 cm dishes at 2 × 10⁶ cells/well (2 mL/well) and cultured for 10 days. Half the medium was replaced with fresh complete medium on day 3, 6, 8. On day 10, CD11c⁺ BMDCs were enriched.

### Cell Survival Assay

Enriched CD11c⁺ BMDCs were seeded into 96-well plates at 5×10⁴ cells/well in 100 μL complete RPMI 1640 medium, and stabilized for 6 h in a 37 °C, 5% CO₂ incubator. Cells were treated with drug combinations at the following concentrations: 20 ng/mL TNF-α (Cell Sciences, CRT192C), 100 nM SMAC mimetics (MCE, HY-12600), 20 μM z-VAD-fmk (MCE, HY-16658), 100 ng/mL LPS (Sigma-Aldrich, L2880/L2630). After 10 h of treatment, cell survival was detected with CellTiter-Glo® Luminescent Cell Viability Assay kit (Promega, G757) according to the manufacturer: 100 μL reagent was added to each well, shaken for 2 min, incubated at room temperature in the dark for 10 min, and luminescence intensity was recorded with a microplate luminometer (Thermo Scientific). Cell survival rate = (luminescence of treatment group / luminescence of control group) × 100%.

### Immunoblotting (Western Blot)

CD11c-enriched BMDCs were harvested post-treatment, washed twice with ice-cold PBS, and finally lysed to 1× SDS sample buffer supplemented with 100 mM DTT (Beyotime, ST043) after quantification by BCA assay (Beyotime, P0010S). Lysates were denatured at 95 ℃ for 8 min, centrifuged at 13,200 rpm (16,000 × g) for 10 min at 4 °C to remove debris, and separated by 10% SDS-PAGE, and transferred to PVDF membranes (Millipore, IPVH00010) at 110 V for 1.5 h at 4 °C. Membranes were blocked with 5% non-fat milk/TBST, incubated overnight at 4 °C with primary antibodies (rabbit anti-RIPK1, CST, 3493P, 1:1000; mouse anti-β-actin, Sigma-Aldrich, A3854-200UL, 1:5000, loading control), followed by HRP-conjugated secondary antibodies (Jackson ImmunoResearch, 1:5000) for 1 h at room temperature. Target proteins were detected with ECL substrate (Thermo Scientific) using a chemiluminescence imaging system (Tanon).

### Immunofluorescence Staining and Confocal Microscopy

HEPG2 cells were seeded onto 13 mm sterile glass coverslips (Thermo, 174950) in 10 cm dish, and cultured overnight in DMEM (Gibco, C11995500BT) supplemented with 10% FBS (Bioind, 04-001-1A), 1% penicillin/streptomycin (Gibco, 15140122) at 37 °C with 5% CO₂. Cells were washed twice with ice-cold PBS, fixed with 4% PFA (Servicebio, G1101) for 15 min at room temperature, then washed three times with PBS. Non-specific binding was blocked with blocking buffer (Beyotime, P0102) for 1 h at room temperature. Cells were incubated with indicated mouse serum samples in blocking buffer at 4 °C overnight, washed three times with PBS, then incubated with Rhodamine (TRITC) AffiniPure® Goat Anti-Mouse IgG (H+L) antibodies (Jackson ImmunoResearch, AB_2338478) for 1 h at room temperature in the dark. After three PBS washes, coverslips were mounted with anti-fade medium containing DAPI (Beyotime, P0131) for simultaneous nuclear staining and anti-fade mounting. Images were acquired using an Olympus FV3000 laser-scanning confocal microscope (405 nm for DAPI, 488 nm for FITC, 555 nm for TRITC) at 1024×1024 resolution. ≥3 random fields were captured per sample, and images were analyzed with ImageJ.

### Flow Cytometry Analysis

Single-cell suspensions were prepared from thymus, spleens and lymph nodes of 6–8-week-old mice euthanized by CO_2_ inhalation. Organs were smashed through 2-mL syringe plungers and filtered with 70-μm strainers, and centrifuged at 4000 rpm for 5 min at 4 °C. Splenocytes were treated with AKC lysis buffer for 5 min at room temperature to remove red blood cells and centrifuged to obtain pellets.

For staining: cells were resuspended in 100 μL staining buffer (PBS with 3%BSA, 1mM EDTA, 0.1%NaN_3_), incubated with surface fluorescence-conjugated primary antibodies (CD45, B220, CD3, CD4, CD11b, Ly-6G from BD; CD8a, CD11c, CD317, CD86 from BioLegend; CD19, CD44, CD62L, MHC-II, CD80 from eBioscience; CD40 from Thermo) for 30 min on ice in the dark. If intracellular cytokine staining was requested, cells were washed, fixed and permeabilized with the Foxp3/Transcription Factor Staining Buffer Set (eBioscience, 00-5523-00), and then incubated with fluorescence-conjugated antibodies (IFN-γ, TNF-α from BioLegend) for overnight at 4 ℃ in the dark. After staining, cells were washed and resuspended in 300 μL PBS. Samples were acquired on a CytoFlex S flow cytometer (Beckman Coulter) and analyzed with CytExpert 2.6.

### Tumor Models

6–8-week-old mice were used for tumor model establishment. Tumor cell lines (B16-F10, Pan02, MC38) were cultured in DMEM/RPMI 1640 with 10% FBS and 1% penicillin/streptomycin. Before inoculation, cells were resuspended in 37 °C pre-warmed PBS, and kept warm until use. Subcutaneous injection volume was controlled at 30–50 μL/mouse to minimize excessive dispersion.

1. B16-F10 subcutaneous melanoma model: Mice were subcutaneously injected with 5×10⁴ B16-F10 cells into the right forelimb. Tumor growth was monitored for 14/18 days. Tumor volume was measured by water displacement method.
2. Pan02 subcutaneous pancreatic cancer model: Mice were subcutaneously injected with 2.0×10⁶ Pan02 cells into the right forelimb. Tumor growth was monitored for 40 days.
3. B16-F10 lung metastasis model: Mice were intravenously injected with 3×10⁵ B16-F10 cells via tail vein (50 μL/mouse). After 3 weeks, mice were euthanized by CO_2_ inhalation; lungs were harvested and subjected to other experiments.
4. MC38 liver metastasis model: Mice were anesthetized with isoflurane (induction: 3–5%, maintenance: 1.5–2%). A 1 cm abdominal incision was made to expose the spleen, and 1.5×10⁶ MC38 cells (50 μL PBS) were injected into the spleen. After 5 min, the spleen was removed, and the abdomen was sutured. Mice were recovered on a heating pad and monitored for 30 days. Thirty days post-inoculation, mice were euthanized by CO_2_ inhalation and livers were harvested and subjected to other experiments.

### Single-Cell RNA Sequencing (scRNA-Seq) Data Analysis

The GSE212701 scRNA-Seq dataset was downloaded from GEO (https://www.ncbi.nlm.nih.gov/gds) to explore dendritic cell (DC) heterogeneity and cell death-related pathways. All analyses were performed using Seurat (v4.4.0) in R (v4.3.1).

#### 1. Data Preprocessing and Integration

Raw UMI count matrices of GSE212701 were imported into R via Read10X function and converted to Seurat objects in Seurat. Stringent QC was applied: cells were retained with percent.mt ≤ 5%, nCount_RNA 5,000–50,000 and nFeature_RNA 2,500–5,000. After QC, data were normalized, variance-stabilized and scaled using SCTransform, then integrated with IntegrateData to eliminate batch effects. PCA was performed with RunPCA (20 PCs retained) for subsequent dimensionality reduction and clustering, as determined by ElbowPlot.

#### 2. DC Identification and Annotation

Cell types were annotated using established marker genes from the original publication^45^. Unsupervised clustering was performed with FindNeighbors and FindClusters (resolution = 0.8), and marker gene expression was visualized via DotPlot and VlnPlot. Non-DC populations (T cells, monocytes, macrophages, B cells, NK cells) were excluded. DCs were identified by specific markers (HLA-DRA, CD1C, CLEC9A and XCR1, etc.) and extracted for the downstream analysis.

#### 3. Pro-Death Gene Signature and GSVA Scoring

A custom pro-death gene signature was constructed to evaluate the heterogeneity of cell death pathway activation in DCs. Specifically, gene sets related to the positive regulation of apoptosis and necroptosis were retrieved from the Gene Ontology (GO) and Kyoto Encyclopedia of Genes and Genomes (KEGG) databases. Then, Gene Set Variation Analysis (GSVA) was performed on the normalized DC expression matrix to quantify the enrichment level of the pro-death term in each individual DC. This analysis generated a continuous “Death Score” for each cell, where a higher score indicated stronger activation of pro-death signaling pathways.

#### 4. Identification of Differentially Expressed Genes (DEGs) and Gene Set Enrichment Analysis (GSEA)

To functionally characterize the biological processes between DC subpopulations with distinct pro-death activity, the DC population was re-clustered using the Death Score as the sole input feature. Differentially expressed genes (DEGs) between the resulting death-related clusters were identified using the FindAllMarkers, which employs the Wilcoxon rank-sum test.

Gene Set Enrichment Analysis (GSEA) was performed to functionally annotate the biological processes and signaling pathways associated with the DEGs. First, DEGs were ranked by their avg_log2FC values (from highest to lowest). Pre-defined gene sets from the Molecular Signatures Database (MSigDB), including Hallmark, GO and KEGG collections, were used as reference gene sets. The results were visualized with hierarchical tree diagrams (to reveal relationships between gene sets).

### Statistical Analysis

Data presented in this article are representative results of at least three independent experiments. The statistical significance of data was evaluated by Student’s *t*-test and Wilcoxon test, and the statistical calculations were performed with R.

## SUPPLEMENTARY INFORMATION

Supplementary information includes six figures.

## AUTHOR CONTRIBUTIONS

Y.J.O. and H.B.Z. designed the study; Y.J.O. and L.X.W. performed the experiments and analyzed the data; X.X.Z. conducted mouse phenotypic analyses; X.M.Z. assisted with genotyping; L.M assisted with the tumor analysis; X.X.W. supported mouse experiments and technical guidance; L.H. and X.K. contributed to scientific discussion and provided essential resources; Y.J.O., L.X.W. and H.B.Z. wrote the manuscript.

## ACKNOWLEDGEMENTS

This work was supported by the National Key Research and Development Program of China (2022YFA0807300), the National Science and Technology Major Project for Chronic Non-communicable Diseases (2024ZD0531300), the National Natural Science Foundation of China (32270803, 32300630, 32400626), the Shanghai Excellent Academic/Technical Leader Program (22XD1404500), the Shanghai Science and Technology Commission (23141902800), and the China Postdoctoral Science Foundation (2020M671261, GZB20240789, 394142). Support was also provided by the Shanghai Postdoctoral Excellence Program (2024713, 2024715) and the Shanghai Municipal Science and Technology Major Project.

## CONFLIC OF INTEREST

The authors declare no conflict of interest.

---

**Figure S1.**
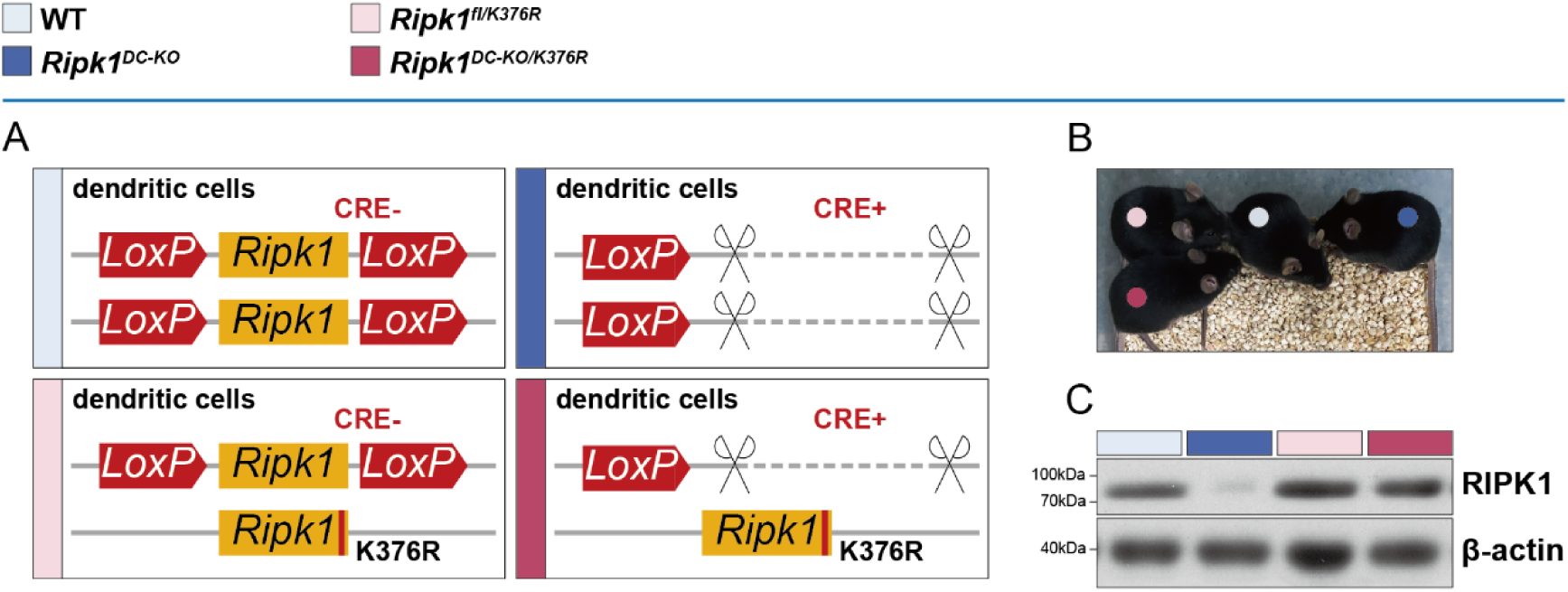
Mouse construction and verification. **(A)** Graphical illustrations of mouse construction strategies of *Ripk1^DC-KO^*, *Ripk1^fl/K376R^* and *Ripk1^DC-KO/K376R^*. **(B)** Representative images of 6-week-old WT, *Ripk1^DC-KO^*, *Ripk1^fl/K376R^* and *Ripk1^DC-KO/K376R^* mice. **(C)** Immunoblot of RIPK1 expressions in splenic CD11c^+^ cells. Spleens were harvested from 8-week-old WT, *Ripk1^DC-KO^*, *Ripk1^fl/K376R^* and *Ripk1^DC-KO/K376R^* mice, and CD11c^+^ cells were enriched using EasySep™ Mouse T Cell Isolation Kit (STEMCELL, 19851).

**Figure S2.**
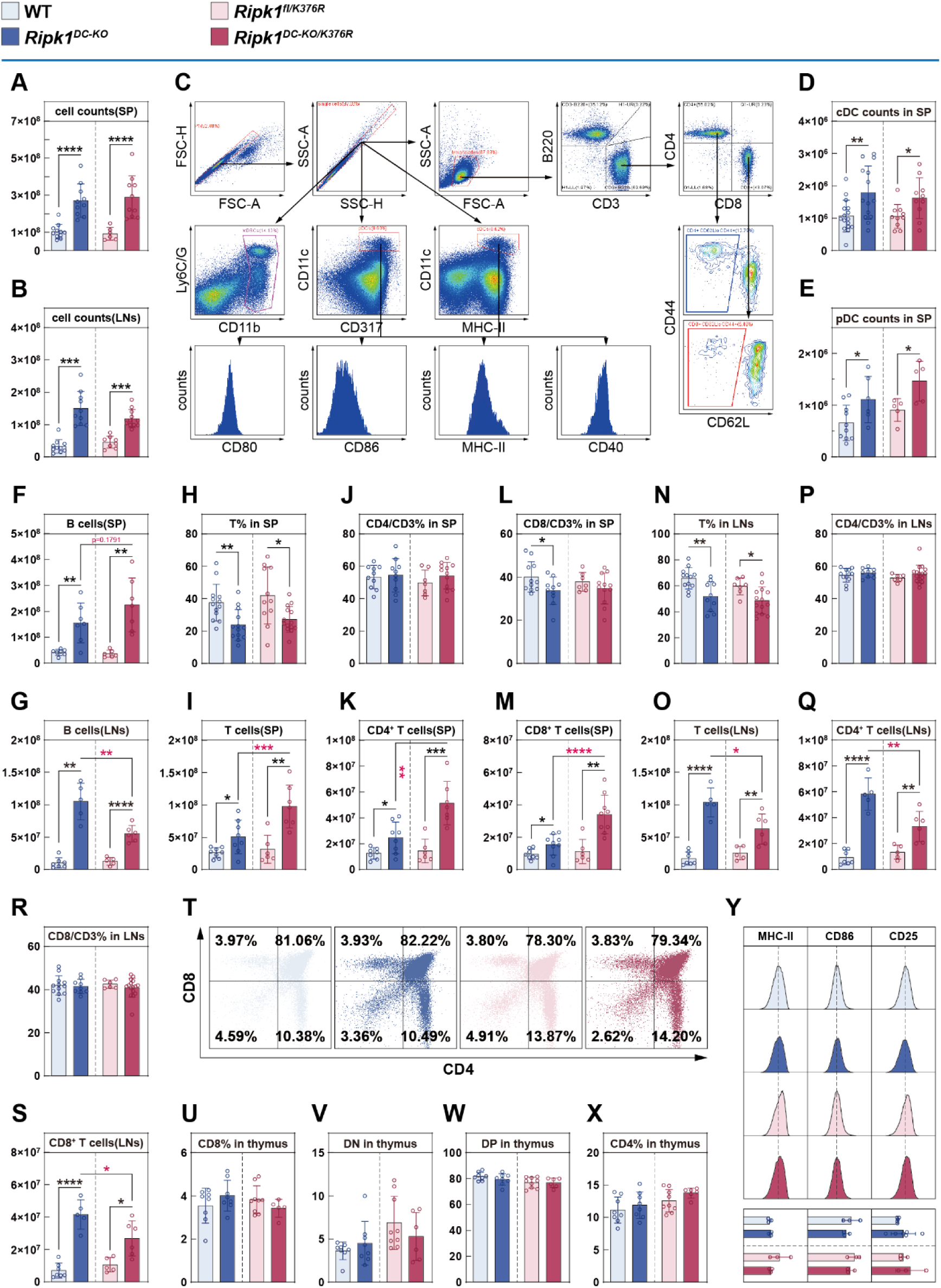
Differences and similarities of lymphocytes between indicated genotypes of mice. **(A, B)** Lymphocyte cell counts of spleens and lymph nodes in 6-week-old WT, *Ripk1^DC-KO^*, *Ripk1^fl/K376R^* and *Ripk1^DC-KO/K376R^* mice (n = 6–11). **(C)** Gating strategy for the identification of indicated immune cell subsets via flow cytometry. **(D, E)** Cell counts of splenic CD11c⁺ MHC-Ⅱ⁺ cDCs and CD11c^lo^ PDCA-1⁺ pDCs in 6-week-old WT, *Ripk1^DC-KO^*, *Ripk1^fl/K376R^*and *Ripk1^DC-KO/K376R^* mice (n = 5–12). **(F, G)** Cell counts of B cells in spleens and lymph nodes in 6-week-old WT, *Ripk1^DC-KO^*, *Ripk1^fl/K376R^* and *Ripk1^DC-KO/K376R^* mice (n = 6–7). **(H–S)** Flow cytometric quantifications and cell counts of total T cells and CD4^+^/CD8^+^ T cell subsets in spleens and lymph nodes in 6-week-old WT, *Ripk1^DC-KO^*, *Ripk1^fl/K376R^* and *Ripk1^DC-KO/K376R^*mice (n = 5–16). **(T–X)** Representative images of flow cytometry (T) and quantifications (U–X) of CD4⁺CD8⁺ double-positive, CD4⁺CD8⁻ single-positive, CD4⁻CD8⁺ single-positive and CD4⁻CD8⁻ double-negative T cell subsets in the thymus in 6-week-old WT, *Ripk1^DC-KO^*, *Ripk1^fl/K376R^* and *Ripk1^DC-KO/K376R^* mice (n = 5–9). **(Y)** Representative images of flow cytometry (up) and quantifications (down) of B cell surface markers MHC-Ⅱ, CD86 and CD25 in 6-week-old WT, *Ripk1^DC-KO^*, *Ripk1^fl/K376R^* and *Ripk1^DC-KO/K376R^*mice (n = 3–6). Data are presented as the mean ± SD from at least three independent experiments, with dots representing individual samples. Statistical significance (*P* values) was determined by a two-tailed unpaired Student’s t-test or Wilcoxon test (*p < 0.05, **p < 0.01, ***p < 0.001, ****p < 0.0001).

**Figure S3.**
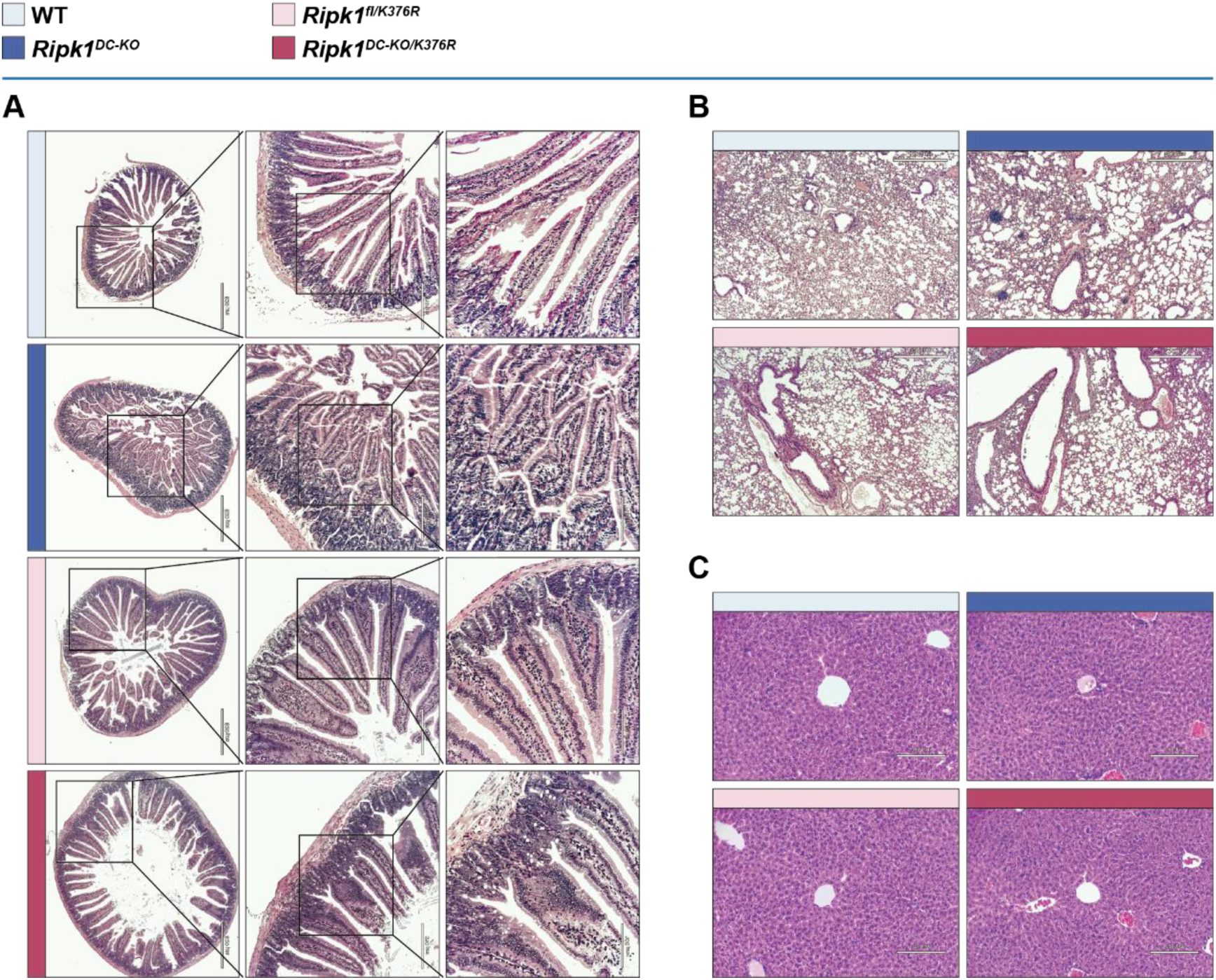
Limited effects of autoimmunity on different tissues. **(A-C)** Representative images of HE staining duodenums (A), lungs (B) and livers (C) in 6-week-old WT, *Ripk1^DC-KO^*, *Ripk1^fl/K376R^*and *Ripk1^DC-KO/K376R^* mice.

**Figure S4.**
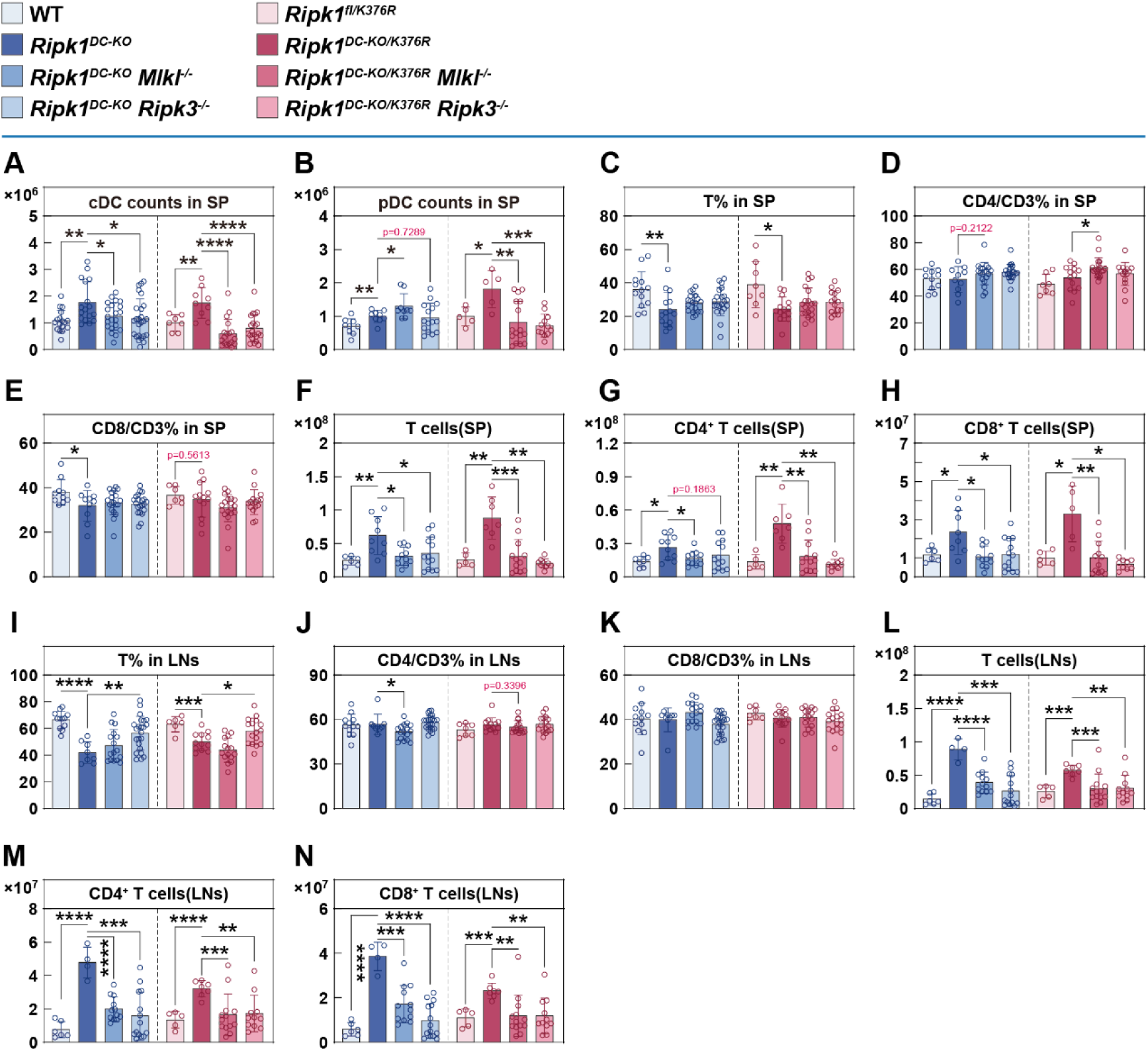
Dendritic cell necroptosis contributes to lymphocyte dysregualtion. **(A, B)** Cell counts of splenic CD11c^+^ MHC-Ⅱ^+^ cDCs (A) and CD11c^lo^ CD317^+^ pDCs (B) in 6-week-old WT, *Ripk1^DC-KO^*, *Ripk1^DC-KO^ Mlkl^-/-^*, *Ripk1^DC-KO^ Ripk3^-/-^*, *Ripk1^fl/K376R^*, *Ripk1^DC-KO/K376R^*, *Ripk1^DC-KO/K376R^ Mlkl^-/-^* and *Ripk1^DC-KO/K376R^ Ripk3^-/-^* mice (n = 5-25). **(C-H)** Flow cytometric quantifications (C-E, n = 7-21) and cell counts (F-H, n = 5-13) of splenic T cells and corresponding CD4^+^/CD8^+^ T cell subsets in 6-week-old WT, *Ripk1^DC-KO^*, *Ripk1^DC-KO^ Mlkl^-/-^*, *Ripk1^DC-KO^ Ripk3^-/-^*, *Ripk1^fl/K376R^*, *Ripk1^DC-KO/K376R^*, *Ripk1^DC-KO/K376R^ Mlkl^-/-^* and *Ripk1^DC-KO/K376R^ Ripk3^-/-^* mice. **(I-N)** Flow cytometric quantifications (I-K, n = 6-23) and cell counts (L-N, n = 4-14) of lymphatic T cells and corresponding CD4^+^/CD8^+^ T cell subsets in 6-week-old WT, *Ripk1^DC-KO^*, *Ripk1^DC-KO^ Mlkl^-/-^*, *Ripk1^DC-KO^ Ripk3^-/-^*, *Ripk1^fl/K376R^*, *Ripk1^DC-KO/K376R^*, *Ripk1^DC-KO/K376R^ Mlkl^-/-^* and *Ripk1^DC-KO/K376R^ Ripk3^-/-^* mice. Data are presented as the mean ± SD from at least three independent experiments, with dots representing individual samples. Statistical significance (*P* values) was determined by a two-tailed unpaired Student’s t-test or Wilcoxon test (*p < 0.05, **p < 0.01, ***p < 0.001, ****p < 0.0001).

**Figure S5.**
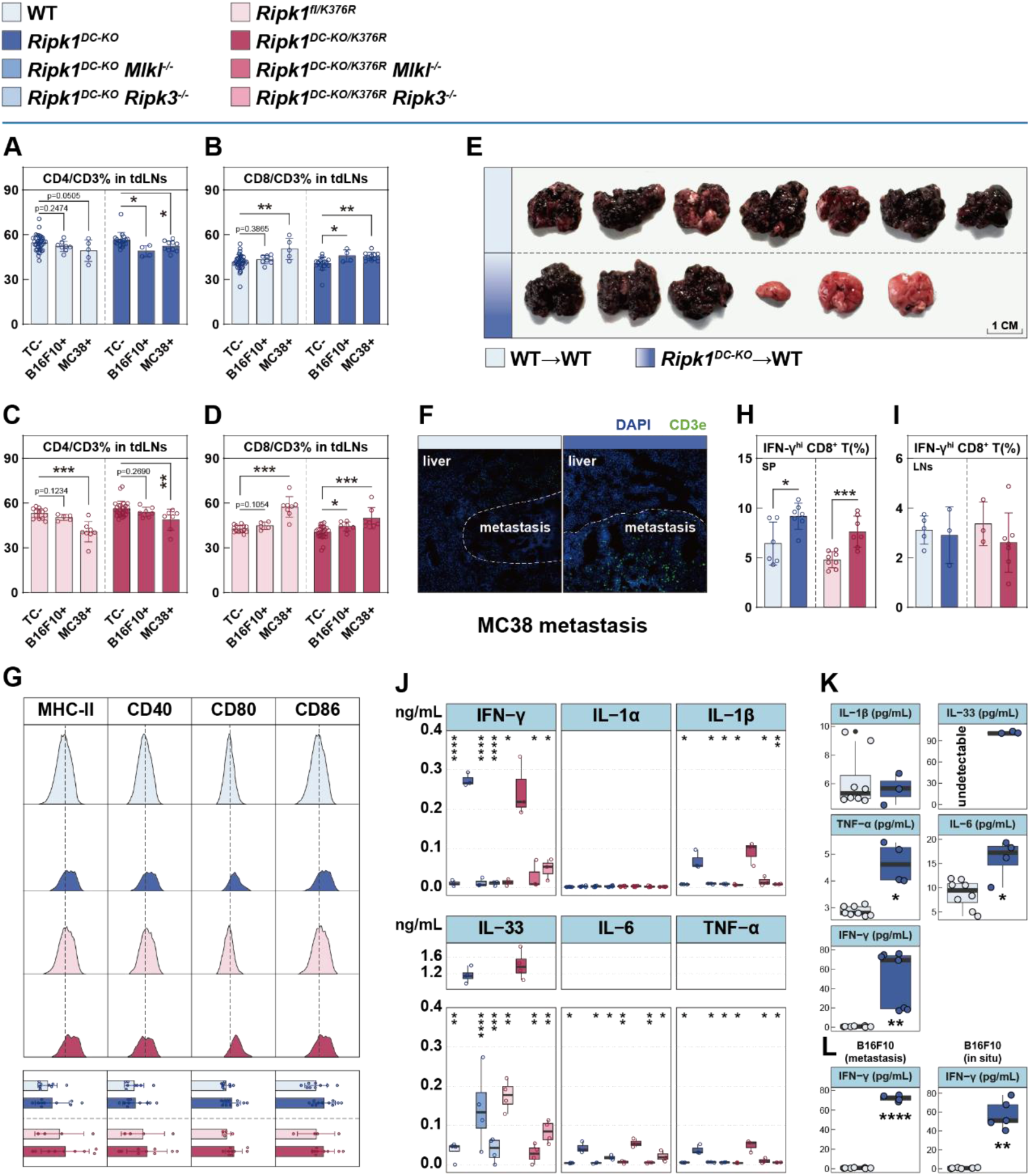
T cell activation contributes to pan-tumor resistance. **(A-D)** Flow cytometric quantifications of CD4^+^ (A) and CD8^+^ (B) T cell subsets in tumor draining lymph nodes in WT, *Ripk1^DC-KO^*, *Ripk1^fl/K376R^* and *Ripk1^DC-KO/K376R^* mice (n = 4-38). **(E)** The image of B16-F10 lung metastatic nodules in T cell transferred mice 21 days after intravenous injection. **(F)** Representative images of immunofluorescence of CD3e and DAPI of MC38 liver metastasis of WT and *Ripk1^DC-KO^* mice. **(G)** Representative images of flow cytometry (up) and quantifications (down) of DC surface markers MHC-Ⅱ, CD40, CD80 and CD86 in 6-week-old WT, *Ripk1^DC-KO^*, *Ripk1^fl/K376R^* and *Ripk1^DC-KO/K376R^* mice (non-tumor-challenged) (n = 4-11). **(H, I)** Flow cytometric quantifications of splenic CD8^+^ IFN-γ^hi^ T cells of 6-week-old WT, *Ripk1^DC-KO^*, *Ripk1^fl/K376R^*and *Ripk1^DC-KO/K376R^* mice (non-tumor-challenged) (n = 3-8). **(J)** ELISA of mouse serum cytokines (IFN-γ, IL-1α, IL-1β, IL-33, IL-6, TNF-α) in 6-week-old WT, *Ripk1^DC-KO^*, *Ripk1^DC-KO^ Mlkl^-/-^*, *Ripk1^DC-KO^ Ripk3^-/-^*, *Ripk1^fl/K376R^*, *Ripk1^DC-KO/K376R^*, *Ripk1^DC-KO/K376R^ Mlkl^-/-^* and *Ripk1^DC-KO/K376R^ Ripk3^-/-^* mice (non-tumor-challenged) (n = 3-6). **(K, L)** ELISA of mouse serum cytokines, including IL-1β, IL-33, TNF-α, IL-6, IFN-γ in WT and *Ripk1^DC-KO^* mice (n = 3-8). Mouse serum was harvested from MC38 challenged mice (K) and B16-F10 challenged mice (L). Data are presented as the mean ± SD from at least three independent experiments, with dots representing individual samples. Statistical significance (*P* values) was determined by a two-tailed unpaired Student’s t-test or Wilcoxon test (*p < 0.05, **p < 0.01, ***p < 0.001, ****p < 0.0001).

**Figure S6.**
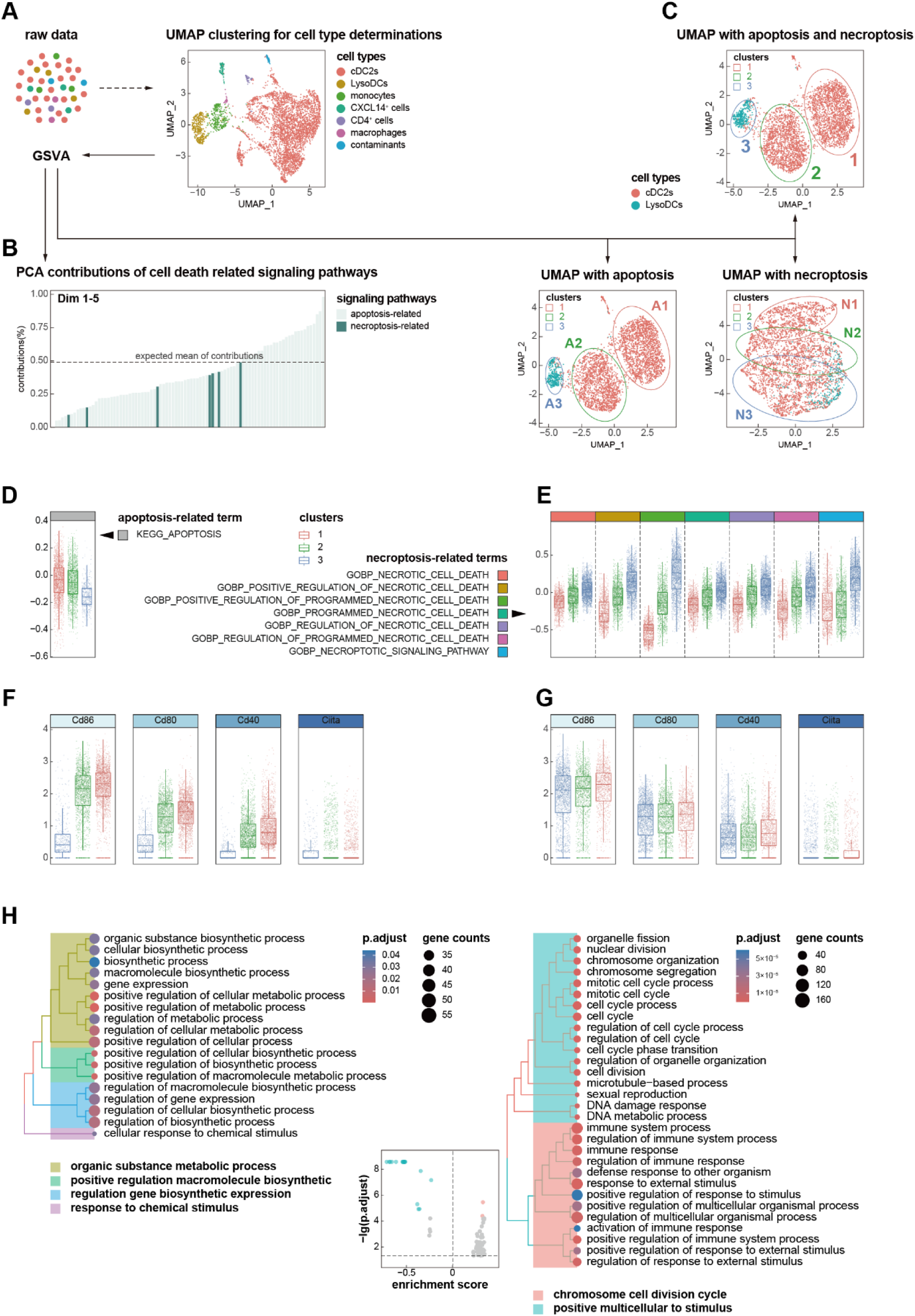
Dendritic cell death has effects on their maturation and activation. **(A)** The GSE212701 dataset was preprocessed for cell type clustering; non-target cells were excluded, and GSVA scores were calculated for positively regulated apoptosis and necroptosis gene sets retrieved from the GO and KEGG databases. **(B)** Principal component analysis (PCA) was performed using cell death-related GSVA scores to evaluate the relative contributions of apoptosis and necroptosis. **(C)** Unsupervised cell clustering was conducted based on combined apoptosis/necroptosis signatures (up), or on each signature individually (down). **(D, E)** Boxplots showing the distribution of apoptosis (D) and necroptosis (E) GSVA scores across PCA-derived cell clusters. **(F, G)** Boxplots depicting the transcriptions of *Cd86*, *Cd80*, *Cd40* and *Ciita* in subsets stratified by apoptosis (F) and necroptosis (G) GSVA scores. **(H)** Gene set enrichment analysis (GSEA) was performed on differentially expressed genes (DEGs) from Cluster 1 vs. 2 and Cluster 1 vs. 3 comparisons; enrichment scores of necroptosis-associated terms are presented.

